## supplemental figures 1-9 for "Proliferative arrest induces neuronal differentiation and innate immune responses in normal and Creutzfeldt-Jakob Disease agent (CJ) infected rat septal neurons"

**SUPPLEMENTAL FILES**

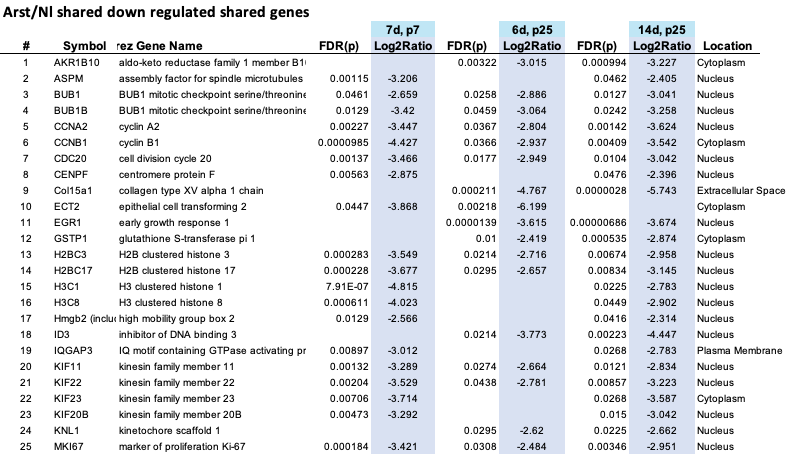

**S1 Table1:** **Top 25 genes down regulated in each of 3 independent samples from different passage of Nl cells,** i.e., Arst/Nl for 7 days from passage 7, at 6 days from p 25, and at 14 days from p25. All genes are downregulated in at least 2 of the 3 samples (see text) with comparable fold changes (Log_2_Ratio columns).

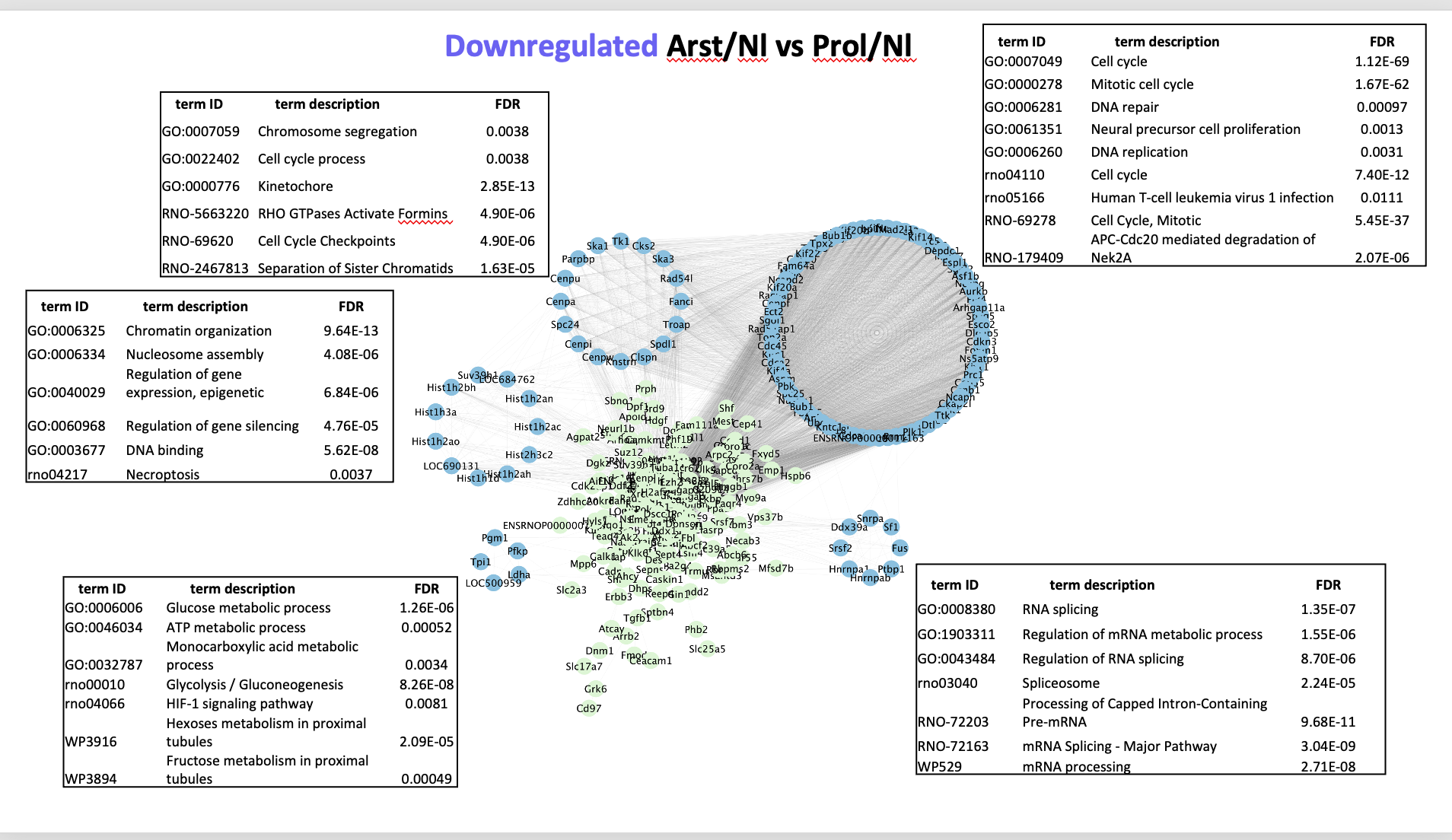

**S2 Fig.: Arts/Nl vs Prol/Nl: Interaction network analysis of differentially downregulated genes.** Interaction network built using STRING knowledgebase (v. 12.0). Network was imported into Cytoscape (v. 3.10.2) and clustering analysis was performed with MCODE (v. 2.0.3) with degree cutoff 2, node density cutoff 0.1, K-core 2, and maximum depth 100. Resulting clusters were functionally annotated on STRING database.

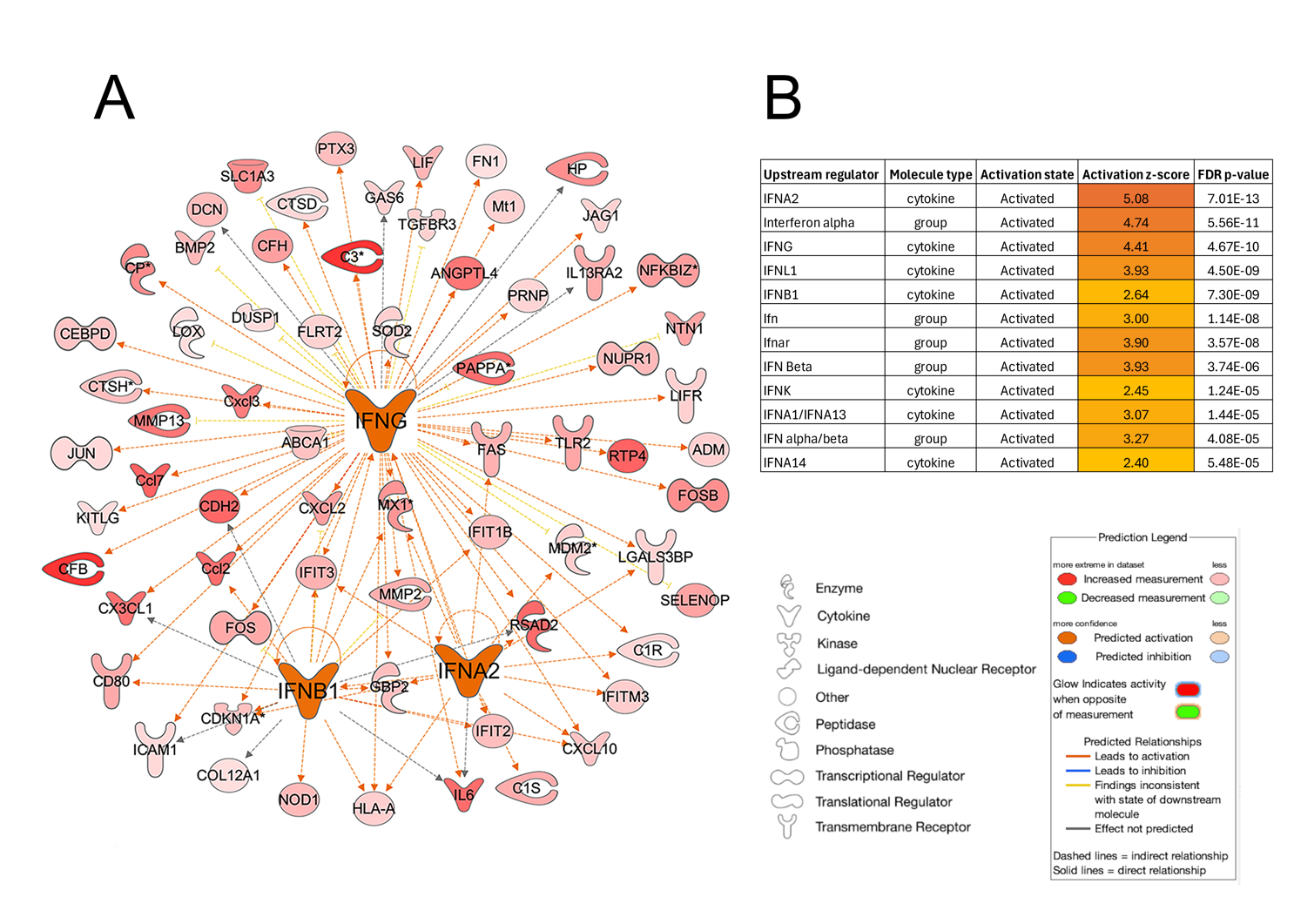

**S3 Fig.: Ingenuity Pathway Analysis for IFN in Nl neurons**. Upstream regulators network for IFNA2, IFNB1, and IFNG (A), and results from upstream regulators prediction and activation status. Intensity of red color in molecules indicates expression levels as shown in the legend. B) Table with prediction confidence and activation z-score value for each molecule.

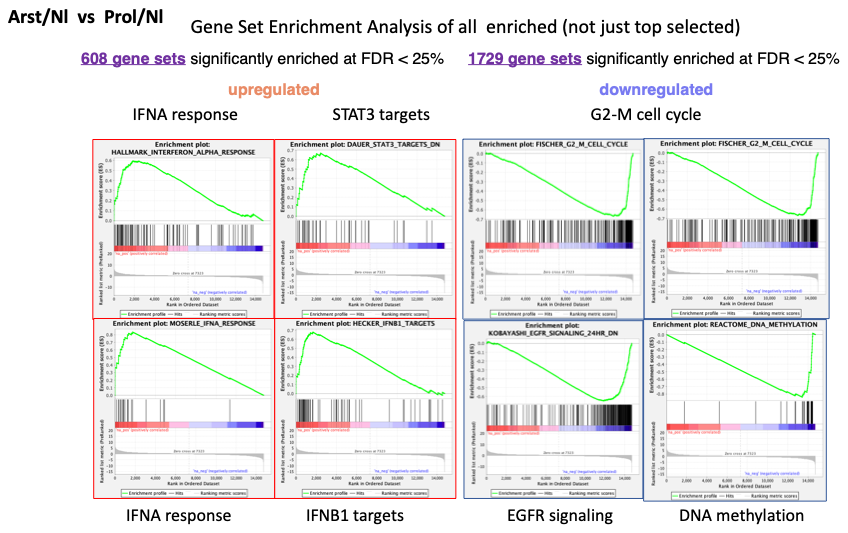

**S4 Fig.:** **Gene set enrichment analysis (GSEA) plots.** Shows enrichment of molecular signatures from two group comparison between Arst/Nl vs Prol/Nl groups.

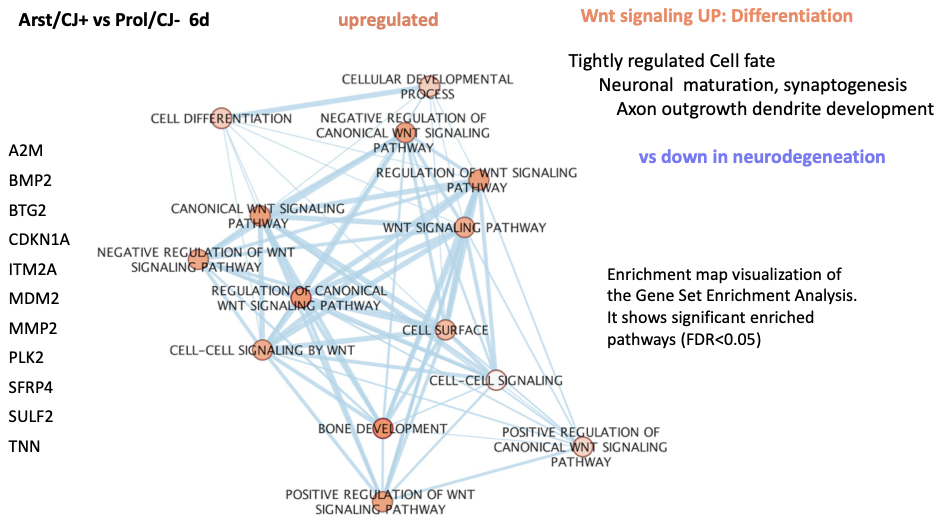

**S5 Fig.: Example of Wnt signaling in arrested CJ+ cells.** Overrepresentation analysis was done on g:Profiler web server and enriched pathways (FDR p<0.05) were clustered using Enrichment map application on Cytoscape. Gene list to the left shows common upregulated genes across those enriched pathways.

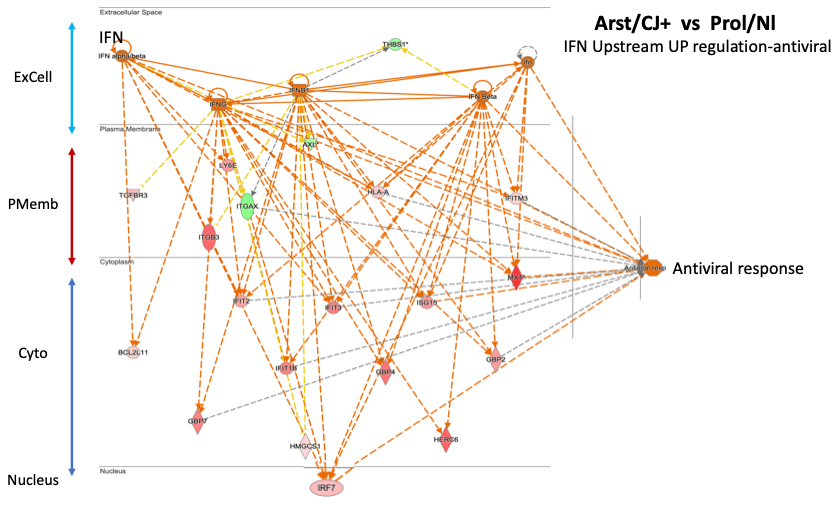
**S6 Fig.:** **CJ+ cells: Ingenuity pathway analysis network**. Overlap between differentially expressed genes and anti-viral response induced by IFN. Activation is indicated by orange lines, and cellular location of the response genes are also shown. In CJ+ cells the IFNs are not extracellular in origin but made by the cell since no other cells are present to produce activating cDNAs. From RT/qPCR direct experimental results in CJ+ cells IFNs were far higher than in uninfected cells indicating above transcripts were enhanced and due to recrudescence of the infectious agent

**IPA path comparisons: Arst vs Prolif**

1) Arst/Nl	vs	Prol/Nl 2) Arst/CJ+ vs	Prol/Nl 3) Arst/CJ+ vs	Prol/ CJ-

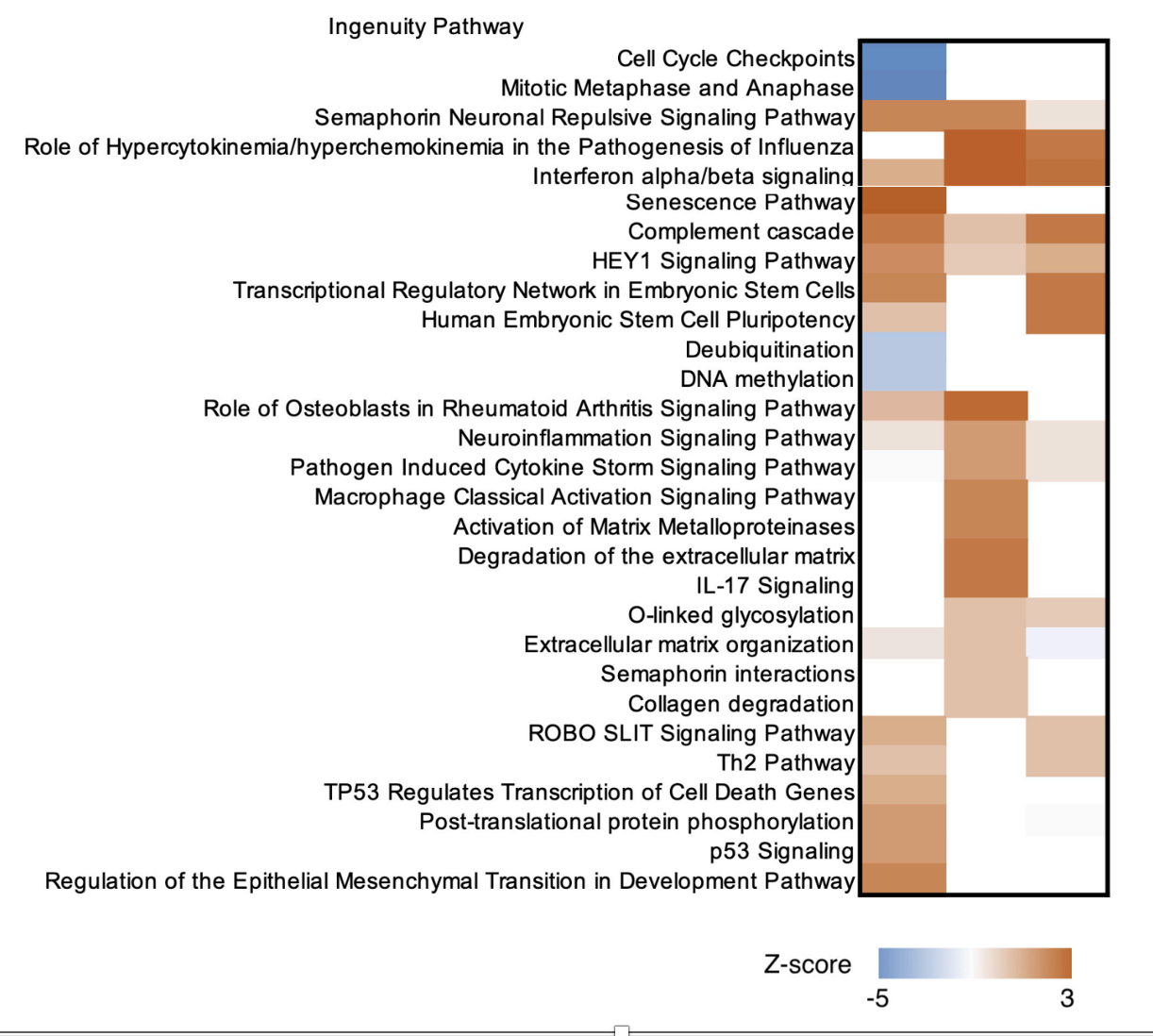

Ar-Nl/Nl CJ+/Nl CJ+/CJ-

**S7 Fig.: Ingenuity pathway analysis of three different comparisons based on pathway activation status (z-score).** Note the difference for the CJ+/CJ- comparison lane 3 with Arst/Nl in lane 1 and CJ+/Nl showing underlying differences in CJ- brought out by arrest. Blue color indicates a negative z-score (inhibition), while orange color indicates a positive z-score and activation.

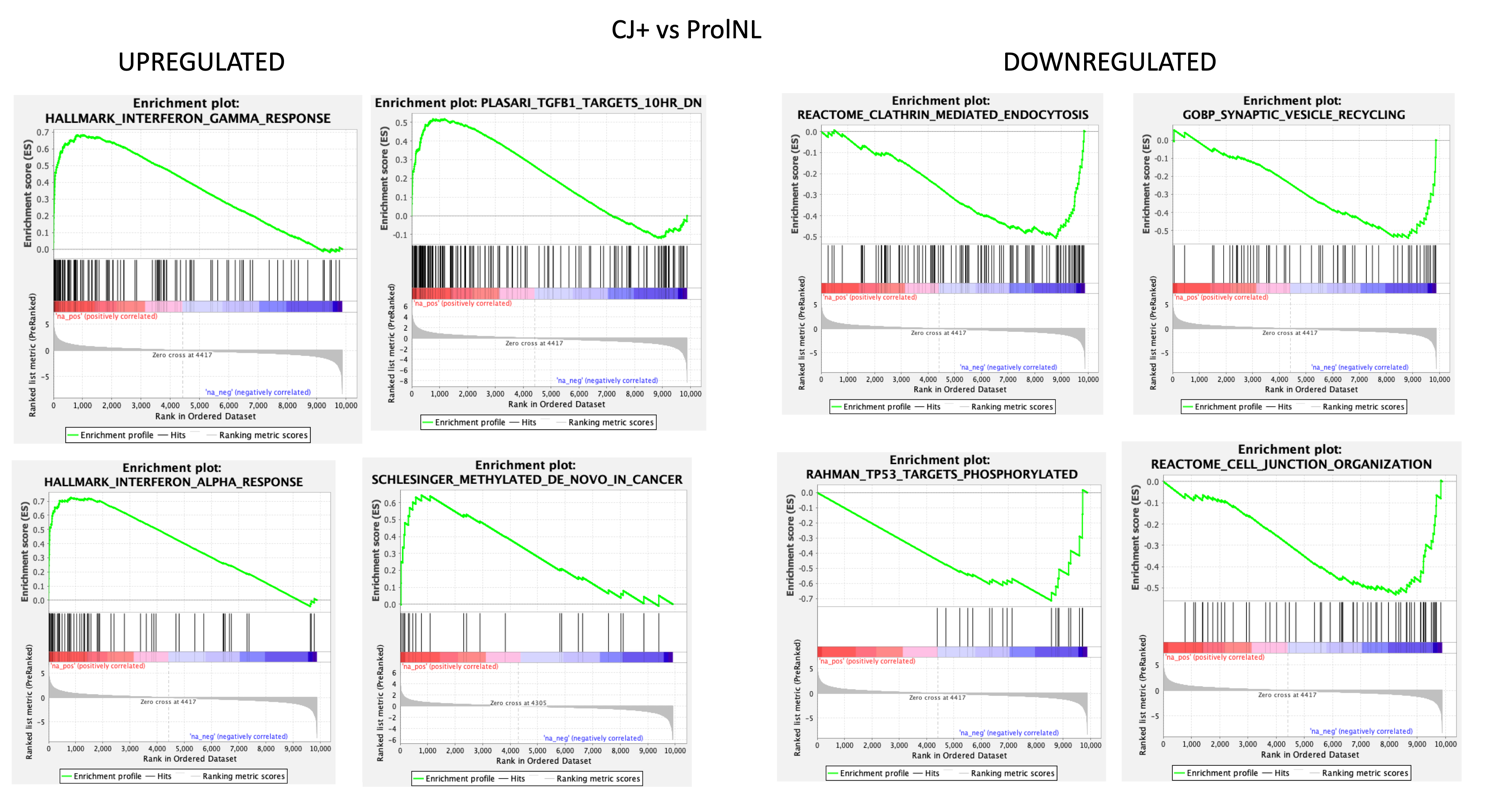

**S8 Fig.: Enrichment plots from Gene Set Enrichment Analysis (GSEA)**. Shows enriched molecular signatures comparing CJ+ versus Prol/Nl samples. Compare S4 above for gene enrichment analyses of Arst/Nl versus Prol/Nl.

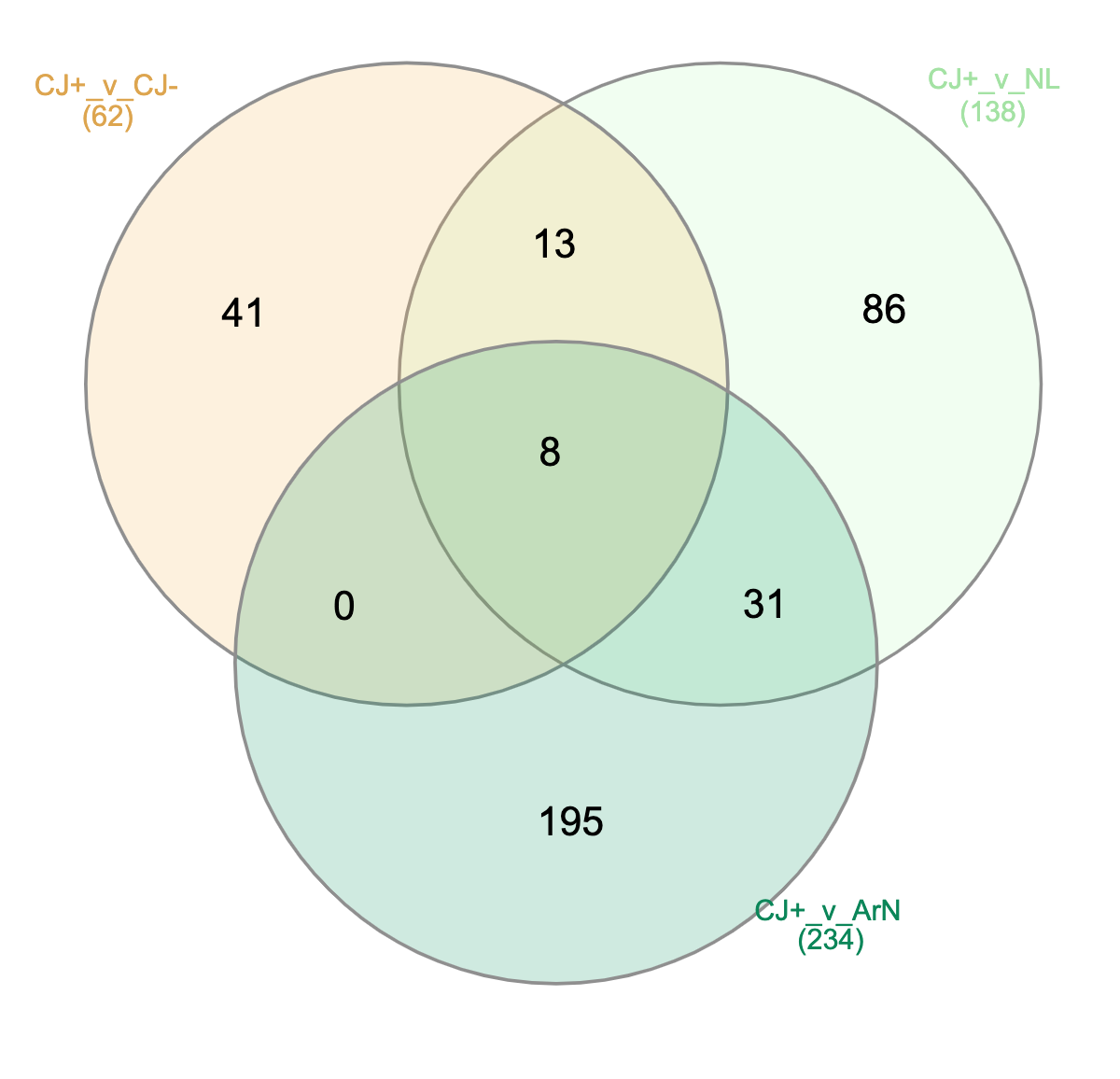

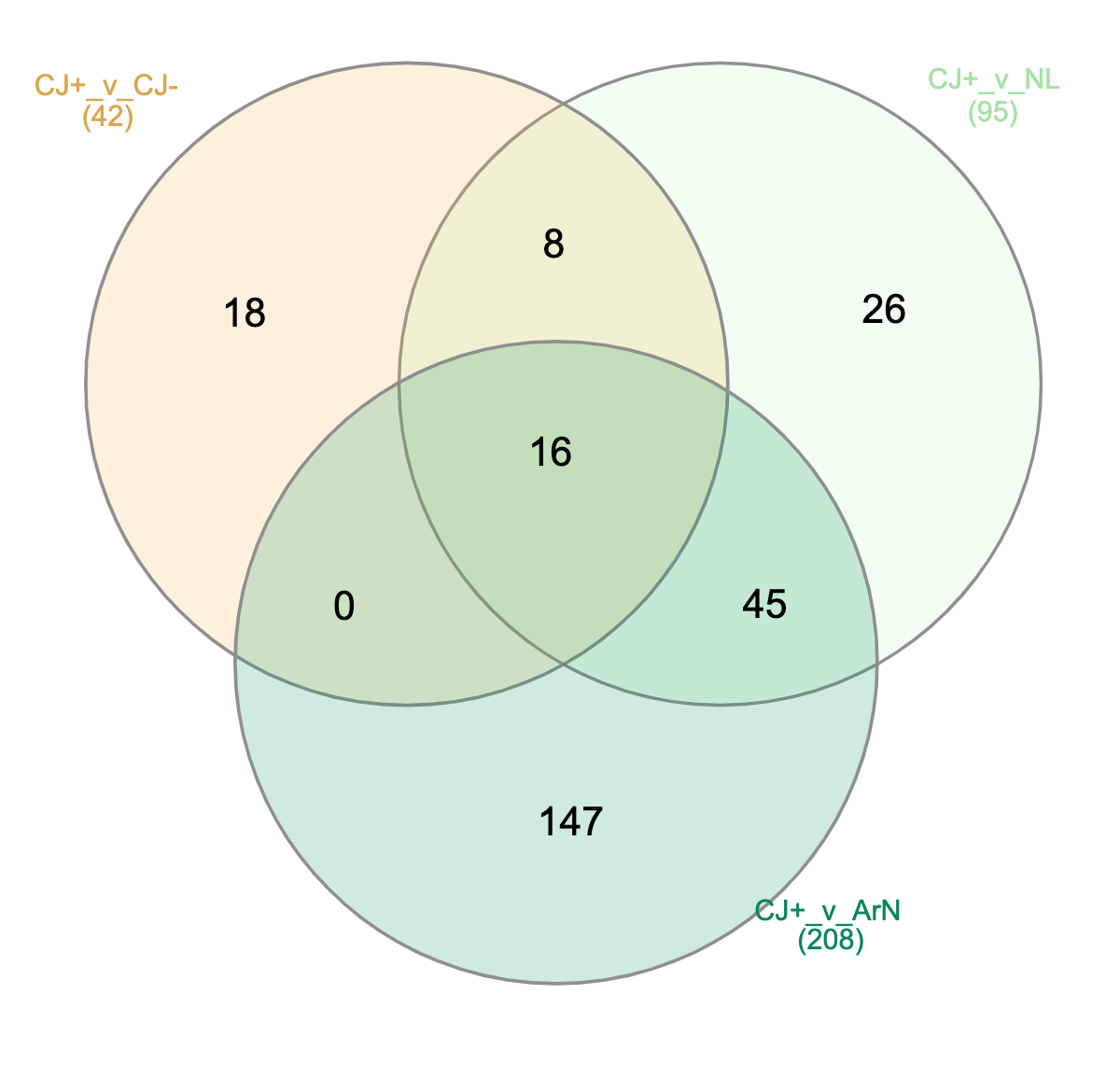

A. B

**S9 Fig.: Venn diagram of unique 196 upregulated CJ+ unique genes** (A) and 146 downregulated genes. B shows 146 CJ+ unique down regulated genes. Please refer to the supplement S10 for complete list.

**S10 excel:** **Complete lists of unique up and down regulated genes in CJ+ versus Prol/CJ- (pink), Prol/Nl (light green) and Arst/Nl (dark green).** Note the minimal differences in Prol/CJ- versus Prol/Nl comparisons and abundance of unique differences by arrest as seen in Arst/CJ+ and Arst/Nl (dark green columns).
